# Depressiveness and Neuroticism in Bartonella Seropositive and Seronegative Subjects – Preregistered Case-Controls Study

**DOI:** 10.1101/275800

**Authors:** Jaroslav Flegr, Marek Preiss, Pavla Balatova

**Author notes:** **Correspondence:** Jaroslav Flegr.

## Abstract

Several recent studies have demonstrated the association of cat-related injuries with major depression and with depressiveness in the general population. It was suggested that cat-scratch disease, the infection with the bacterium *Bartonella henselae*, can be responsible for the observed association. However, no direct evidence for the role of the *Bartonella* infection in this association has been published until now. In this preregistered case-controls study performed on 250 healthy subjects tested earlier for the presence of anti-*Toxoplasma* IgG antibodies, we searched for the positive association between presence of anamnestic anti-*Bartonella* IgG antibodies and depressiveness measured with Beck II inventory, depression subscale of neuroticism measured with N-70 questionnaire, and self-reported health problems. We found that that *Bartonella*-seropositivity was positively correlated with Beck depression only in *Toxoplasma*-seronegative men and negatively correlated with health in *Toxoplasma*-seronegative women. *Bartonella* seropositivity expressed protective effects against *Toxoplasma* seropositivity-associated increased neuroticism in men while *Toxoplasma*-seropositivity expressed protective effects against *Bartonella* seropositivity-associated health problems in women. A comparison of the patterns of association of mental and physical health problems with *Bartonella* seropositivity and with reported cat-related injury suggests that different factor, possibly infection with different pathogen transmitted by cat related-injuries than the *Bartonella henselae*, is responsible for the observed association of cat related-injuries with depressiveness and major depression. The existence of complex interactions between *Bartonella* seropositivity, *Toxoplasma* seropositivity, and sex also suggest that the effect of symbionts on the host’s phenotype must by always studied in the context of other infections, and separately for men and women.

## 1. Introduction

Recently, three independent studies have shown that cat-related injuries (Hanauer et al., 2013), and namely scratching by a cat (Flegr and Hodny, 2016; Flegr and Vedralova, 2017), is statistically associated with suffering from major depression and also with increased depressiveness as measured by the Beck Depression Inventory in non-clinical populations. Based on these observations we suggested that the cat-scratch disease, the infection with gram-negative bacteria of the genus *Bartonella*, can be responsible for this association. These bacteria are present in blood and other bodily fluids of about 50 % of cats in many parts of world (Massei et al., 2005). The infection causes relatively common cat-scratch disease as well as several other less frequent but more serious diseases such as infectious endocarditis, bacillary angiomatosis, and Oroya fever (Prutsky et al., 2013; Breitschwerdt, 2014). *Bartonella* is transmitted from cats to humans by flea bites and by cat scratches, mostly by contamination of scars by bacteria-containing fleas’ feces. It can probably also be transmitted by cat bites, and possibly also by ticks. Cats are most often infected with *Bartonella henselae*; however, several other species of the genus *Bartonella* are present also in cats and other hosts (Angelakis and Raoult, 2014). The incidence of recognized (or reported) disorder in the USA was only 9.3 per 100,000 inhabitants (Jackson et al., 1993), however, the seroprevalence of bartonellosis in human population is 5-30 %. Infected subjects usually develop skin lesions and unilateral lymphadenitis in the lymph-draining region of the site of injury. Patients can suffer from low-grade fever, aching, malaise, headache, or splenomegaly, distortion of vision, anorexia, abdominal pain, severe liver and spleen tissue abnormalities, sore throat, and conjunctivitis for about 3 to 10 days after the infection. Swollen lymph nodes are typical and take weeks to months to subside. Typically, the disease is self-limiting and patients do not require any treatment. Sometimes, however, the lymphadenopathy persists several months, and more serious sequels, including neuroretinitis, encephalopathy, or osteomyelitis, can occur. Often, neurological symptoms of the infection develop, including severe headache, acute confusion, seizures, ataxia, tremors, and focal neurological deficits (Perkins et al., 1992; Breitschwerdt et al., 2008; Rondet et al., 2012). Some patients also express fatigues, memory loss, and depression (Breitschwerdt et al., 2008). No effective method of treatment of cat-scratch disease is currently available (Prutsky et al., 2013).

In the present preregistered case-control study, we searched for direct evidences for the association between of *Bartonella* seropositivity and mental health, namely depressiveness and neuroticism, and physical health in the nonclinical population of 250 subjects. These subjects, mostly former university students of biology, already participated in various studies on the effects of latent infection with protozoan parasite *Toxoplasma* on human health, behavior, and personality. It was shown recently that the *Toxoplasma* infection not only influences the physical and mental health of the infected subjects but also modifies the response of the host organism on the effect of other pathogen, namely spirochete *Borrelia burgdorferi*. The large-scale internet study showed that the increased depressiveness was observed only in *Toxoplasma* co-infected, *Borrelia*-infected subjects (Flegr and Horacek, 2017). To enable the detection of a similar interaction between *Bartonella* and *Toxoplasma* and to take advantage of the fact that all subjects had been tested for the presence of anti-*Toxoplasma* antibodies, we included approximately the same number of *Toxoplasma* seropositive and seronegative subjects into our sample and included the factor *Toxoplasma*-seropositivity into all statistical models.

## 2. Material and Methods

This is a preregistered case-control study Flegr, J. (2017, August 30) “Effects of bartonellosis on depression and health”, osf.io/agvbj. The experimental setup, four hypotheses to be tested and statistical tests for testing these particular hypotheses, as well as the most important follow up tests had been registered before the sera were examined for the presence of anti-*Bartonella* IgG antibodies.

### 2.1. Participants

Data were collected during several projects studying the effects of latent toxoplasmosis on human behavior, personality and health that were performed at the Faculty of Science, Charles University over the past 10 years. During these projects, about 3000 subjects completed various questionnaires in an electronic or paper form and provided a sample of blood for serological testing for *Toxoplasma* and other pathogens. These samples are kept frozen in - 18°C. For the purpose of the current project, we selected a sample of 250 subjects (we tried to find the same number of *Toxoplasma*-seropositive and *Toxoplasma*-seronegative subjects of similar age) who completed an anamnestic questionnaire containing 20 health-related questions, Beck Depression Inventory (BDI-II - clinical scale measuring depression), and N-70 questionnaire (Czech questionnaire measuring seven subscales of neuroticism, including depression). These sera were examined for the presence of anamnestic titres of anti-*Bartonella* antibodies. The study was approved by Institutional Review Board of Faculty of Science, Charles University (No: 2016/16).

### 2.2. Questionnaires

In the anamnestic questionnaire, we asked the responders about the existence and intensity of various health problems. They were asked to subjectively rate the intensity of their allergic, dermatologic, cardiovascular, digestive, metabolic including endocrine, orthopedic, neurological, and psychiatric problems using 7-point Likert scales. Using the same scales, they were asked how frequently they were suffering headache, recurrent pain, other recurrent health problems, being tired, being tired after returning from work or after travelling by train, how frequently they have a common viral diseases, how frequently they have to visit medical doctors (except dentist and except for prevention), how many times they used antibiotics during the past 365 days, how many times they used antibiotics during the past three years, how many times they spent more than one week in a hospital within last year, and how many times they spent more than one week in a hospital within last three years. The participants also completed an electronic form of Czech versions of the Beck Depression Inventory (BDI-II) (Beck et al., 1996) and of N-70 questionnaire (Vacíř, 1973; Flegr et al., 2012). BDI-II is a 21-question multiple-choice (scoring from 0-3 for each item) self-report inventory. BDI-II is one of the worldwide most used psychometric tests for measuring the severity of depression. Higher scores display a higher level of depressive symptomatology. N-70 is a multiple-choice self-report inventory (scoring from 0-3 for each item). The instrument assess several aspects of emotional ability (neuroticism). The N-70 questionnaire is constructed for the assessment of 7 clinical areas - anxiety, depression, phobia, hysteria, hypochondria, vegetative lability (the presence of various psychosomatic symptoms), and psychastenia. The original purpose of this method was to detect individuals who can be too sensitive for military operations (Vacíř, 1973). Scores in each subscale could range from 0-30. The total N-70 score is a sum from all clusters. Higher scores display higher levels of neuroticism. During the preparation of the manuscript, in December 2017, we also sent emails to the participants of the study, asking them about sustained animal-related injuries and taking antidepressants and anxiolytics. Two hundred thirteen subjects provided the requested information.

### 2.3. Serological tests

The complement-fixation test (CFT), which determines the overall levels of IgM and IgG antibodies of particular specificity, and Enzyme-Linked Immunosorbent Assays (ELISA) (IgG ELISA: SEVAC, Prague) were used to detect the *Toxoplasma* infection status of the subjects. ELISA assay cut-point values were established using positive and negative standards according to the manufacturer’s instructions. In CFT, the titre of antibodies against *Toxoplasma* in sera was measured in dilutions between 1:4 and 1:1024. The subjects with CFT titres between 1:8 and 1:128 were considered *Toxoplasma*-seropositive. Only subjects with clearly positive or negative results of CFT and IgG ELISA tests were diagnosed as *Toxoplasma*-seropositive or *Toxoplasma*-seronegative, whilst subjects with different results of these tests, or ambiguous results, were retested or excluded from the study. Frozen samples of sera after 1: 64 dilution were also tested for the presence of anamnestic titres of anti-*Bartonella* IgG antibodies using Indirect Immunofluorescent Assay (FOCUS Diagnostics, U.S.A.). All tests were performed in the reference laboratories of the National Institute of Public Health, Prague.

### 2.4. Statistics

Statistical testing was performed using the programs Statistica v. 10.0 and SPSS v. 21. Before the analyses, the health index was computed for each participant as the average of Z-scores of 20 health-related variables of the anamnestic questionnaire. Participants with intensive or frequent health problems had high (positive) values, while healthy participants had low (negative) values of this index. A logistic regression was used for searching for effects of the predictors age, sex, size of settlement in which a subject spend its childhood, and *Toxoplasma* (*Bartonella*) seropositivity on the probability of being *Bartonella* (*Toxoplasma*) seropositive. ANCOVA tests with a BDI-II score (or individual N-70 subscales scores, or health score) as an output variable were used to search for the effects of the predictors *Bartonella*, age, sex, size of place of living in childhood, *Toxoplasma*, and the sex-*Bartonella, Toxoplasma*-infection-*Bartonella* and sex-*Toxoplasma*-*Bartonella* interactions. The effects of these predictors on six (preregistered) or all seven N-70 variables were studied with MANCOVA. In parallel, the effects of *Bartonella* and *Toxoplasma* seropositivities were also measured with nonparametric partial Kendall test that allow controlling for one confounding variable, here age (Siegel and Castellan, 1988). This method was also used for searching for the effects of seropositivities on responses to a particular 21 questions of BDI-II and particular health problems in the exploratory part of the study. Ordinal regressions were used with 20 different health-related variables (rated on the 7-points scales) as the dependent variables and sex, age, size of settlements where the responders spent their childhood, *Bartonella* seropositivity, *Toxoplasma* seropositivity, and *Bartonella*-sex, *Bartonella*-*Toxoplasma*, and *Bartonella*-*Toxoplasma*-sex interactions as the independent variables. The false discovery rate (preset to 0.20) was controlled with the Benjamini-Hochberg procedure (Benjamini and Hochberg, 1995).

Terminological notes: we abbreviated “*Toxoplasma* seropositivity” to “*Toxoplasma*”, and “*Bartonella* seropositivity” to “*Bartonella*” in the description of our statistical models. For example, 3-way interaction *Toxoplasma* seropositivity-*Bartonella* seropositivity-sex interaction is described in a more condensed way as *Toxoplasma*-*Bartonella*-sex interaction. Also, the statistical relations between (formally) dependent and (formally) independent variables are called “effects” in the Result section of the paper, despite the real causality relation between the these variables can be different or even non-existing (as expanded upon in the Discussion).

### 2.5. Differences between preregistered and realized protocol

We had to substitute the preregistered ANCOVA test for depression subscale from N-70 with non-parametric tests as this variable failed in the Levene’s test of equity of errors. In addition to the preregistered tests with two 2-way interactions (sex-*Bartonella, Toxoplasma*-infection-*Bartonella*) we always additionally analyzed the full model with all three 2-way interactions (sex-*Bartonella, Toxoplasma*-infection-*Bartonella*, and sex-*Toxoplasma*) to show the robustness of our results and to follow recommendation of many statisticians concerning the necessity to analyze full models. However, all inferences have been done exclusively on the basis of results of the preregistered tests. We always used more conservative two-sided, instead of the preregistered one-sided variant of tests because the observed effects of the *Bartonella*-seropositivity were in the opposite directions in men and women.

### 2.6. Data availability

All data are available at: https://doi.org/10.6084/m9.figshare.5852511

## 3. Results

### 3.1. Descriptive statistics

The final set contained data from 92 men and 158 women, see Table 1. No significant differences were observed in the mean age of men (26.7, S.D. = 7.00) and women (26.4, S.D. = 8.9) in the whole set (p = 0.75, t_248_ = -0.316) or between *Bartonella*-or *Toxoplasma*-seropositive and seronegative male or female subjects (all p values > 0.35). Most subjects (31.6% men and 25.0 % women) spent their childhood in Prague, 12.0 % men and 15.2 % women in settlements with less than 1000 inhabitants, 17.4 % men and 13.9 % women in settlements with 1-5 thousands of inhabitants, 26.1 % men and 29.8 % women in settlements with 5-50 thousands of inhabitants, 10.9 % men and 4.4 % women in settlements with 50-100 thousands of inhabitants, 8.7 % men and 5.1 % women in settlements with more than 100 thousands of inhabitants, except Prague. A logistic regression with age, sex, size of settlement in which a subjects spent their childhood as the covariates showed no association between the *Bartonella* seropositivity (dependent variable) and the *Toxoplasma* seropositivity (p = 0.468, O.R. = 1.23, C.I._95_ = 0.71-2.13) and no association between the *Bartonella* seropositivity and the covariates. Analogical analysis with the *Toxoplasma* seropositivity as the dependent variable showed that the probability of being *Toxoplasma*-seropositive decreased with the increasing size of the settlement in which a subject spent his or her childhood (p = 0.004, O.R._range_ = 0.35, C.I._95_ = 0.17-0.71); other associations of *Toxoplasma* seropositivity with covariates were nonsignificant.

**Table 1.** Seroprevalence of bartonellosis in *Toxoplasma*-seronegative and *Toxoplasma*-seropositive subjects.

| Sex | *Toxoplasma* | *Bartonella*-<br>negative | *Bartonella*-<br>positive |
| --- | --- | --- | --- |
| women | negative | 59 | 22 (27.16 %) |
| women | positive | 51 | 26 (33.77 %) |
| men | negative | 37 | 17 (31.48 %) |
| men | positive | 25 | 13 (34.21 %) |

### 3.2. Confirmatory section of the study

Four hypotheses have been preregistered before the start of the present study.

#### 3.2.1. Hypothesis 1

*Subjects who have been diagnosed as Bartonella-seropositive express higher levels of depression measured with N-70 inventory than the Bartonella-seronegative subjects when sex, age, toxoplasmosis and size of place of living are controlled.*

The N-70 inventory was completed by 156 women (30.8 % *Bartonella*-seropositive) and 92 men (32.6 % *Bartonella*-seropositive). The Levene’s test of equity of errors showed that a preregistered ANCOVA test cannot be done (F7,240 = 3.56, p = 0.001); therefore only the nonparametric partial Kendall correlation tests with age as a covariate were performed. Fig. 1 suggests the possible existence of sex-*Bartonella*-*Toxoplasma* interaction. The expected higher depression in *Bartonella*-seropositive subjects was observed only in *Toxoplasma*-seronegative men. However, even here the association was not significant (p = 0.136, partial Kenadall Tau = 0.139, n = 54). *Toxoplasma*-seronegative women showed negative nonsignificant association while *Toxoplasma*-seropositive men showed negative nonsignificant association and *Toxoplasma*-seropositive women no association between *Bartonella* and depression. Nonsignificantly lower depression was observed in *Toxoplasma*-seropositive men (p = 0.063, partial Kenadall Tau = -0.210, n = 38), *Toxoplasma*-seronegative women (p = 0.072, partial Kenadall Tau = -0.136, n = 81), and *Toxoplasma*-seropositive women (p = 0.195, partial Kenadall Tau = -0.102, n = 75) – all in comparison with corresponding *Bartonella*-seronegative subjects.

**Fig. 1.**
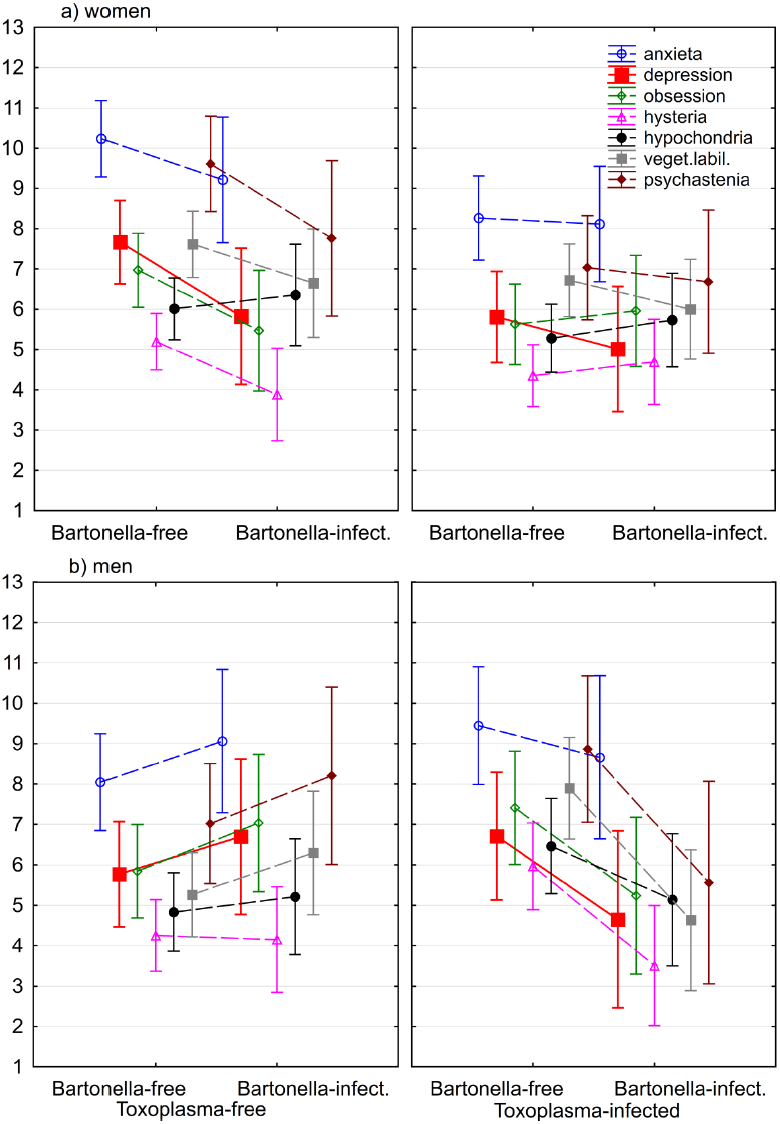
Association of *Bartonella* and *Toxoplasma* seropositivity with seven neuroticism subscales of N-70.

*Markers show the intensity of health problems computed for covariates at their means.*

#### 3.2.2. Hypothesis 2

*Subjects who have been diagnosed as Bartonella-seropositive express higher levels of depression measured with BDI II questionnaire than Bartonella-seronegative subjects when sex, age, toxoplasmosis and size of place of living are controlled.*

The BDI-II questionnaire was completed by 148 women (30.4 % *Bartonella*-seropositive) and 82 men (34.1 % *Bartonella*-seropositive). An ANCOVA test with a BDI-II score as the output variable and *Bartonella*, age, sex, size of place of living in childhood, *Toxoplasma*, and the sex-*Bartonella, Toxoplasma*-*Bartonella* and sex-*Toxoplasma*-*Bartonella* interactions as the independent variables indicated a significant association of depression with the *Toxoplasma*-*Bartonella* interaction (p=0.034, η^2^ = 0.020, two sided test and also with sex-*Toxoplasma*-*Bartonella* interaction (p=0.048, η^2^ = 0.018). Similar results, namely the significant *Toxoplasma*-*Bartonella* interaction (p=0.030, η^2^ = 0.021) and sex-*Toxoplasma*-*Bartonella* interaction (p=0.023, η^2^ = 0.023), also provided the full model with three 2-way and one 3-way interactions. Again, the expected positive *Bartonella*-depression interaction was observed only in *Toxoplasma*-seronegative men. In all three other subpopulations, the *Bartonella*-seropositive subjects expressed lower or the same BDI-II depression scores than the *Bartonella*-seronegative subjects, Fig. 2. Nonparametric partial Kendall correlation tests (two-sided, age controlled) showed significantly higher depression in *Toxoplasma*-seronegative men (p = 0.018, partial Kendall Tau = 0.232, n = 50), non-significantly lower depression in *Toxoplasma*-seropositive men (p = 0.064, partial Kenadall Tau = -0.230, n = 32), and non-significantly higher depression in *Toxoplasma*-seronegative (p = 0.986, partial Kenadall Tau = 0.001, n = 77), and *Toxoplasma*-seropositive women (p = 0.650, partial Kenadall Tau = 0.037, n = 71) – all in comparison with corresponding *Bartonella*-seronegative subjects.

**Fig. 2.**
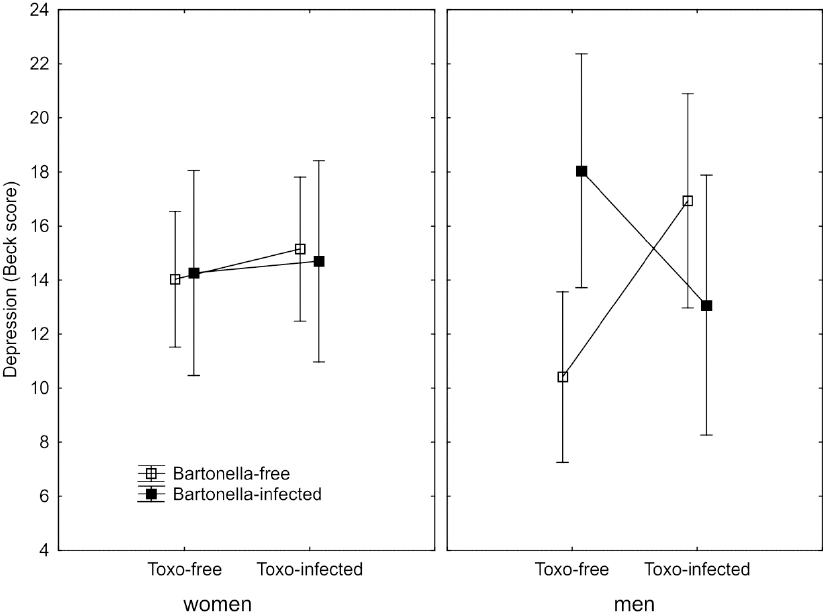
Association of *Bartonella* and *Toxoplasma* seropositivity with BDI-II depression score.

*Squares show depression computed for covariates at their means and spreads 95% confidence intervals.*

#### 3.2.3. Hypothesis 3

*Subjects who have been diagnosed as Bartonella-seropositive report worse health status than Bartonella-seronegative subjects when sex, age, toxoplasmosis and size of place of living is controlled.*

The anamnestic questionnaire was completed by 158 women (30.4 % *Bartonella*-seropositive) and 92 men (32.6 % *Bartonella*-seropositive). An ANCOVA test with the health score as the output variable and *Bartonella*, age, sex, size of childhood home, *Toxoplasma*, and the sex-*Bartonella, Toxoplasma*-*Bartonella* and sex-*Toxoplasma*-*Bartonella* interactions as the independent variables indicated significantly worse health scores in women than in men (p=0.004, η^2^ = 0.034) and also a significant association between health score and *sex*-*Bartonella* interaction (p=0.041, η^2^ = 0.017). Similar results, namely the effects of sex (p=0.005, η^2^ = 0.032), and the *sex*-*Bartonella* interaction (p=0.039, η^2^ = 0.018) also provided the full model with three 2-way and one 3-way interactions. The Fig. 3. shows that *Bartonella* seropositivity had a negative association with health in women and positive association with health in men. Nonparametric partial Kendall correlation tests (age controlled) showed non-significantly better health status of the *Bartonella*-seropositive subjects in *Toxoplasma*-seronegative men (p = 0.646, partial Kenadall Tau = -0.043, n = 54), and in *Toxoplasma*-seropositive positive men (p = 0.080, partial Kenadall Tau = -0.198, n = 38) and also non-significantly worse health status in *Toxoplasma*-seronegative women (p = 0.084, partial Kenadall Tau = 0.131, n = 81), and in *Toxoplasma*-seropositive women (p = 0.428, partial Kenadall Tau = 0.062, n = 77) – all in comparison with corresponding *Bartonella*-seronegative subjects.

**Fig. 3.**
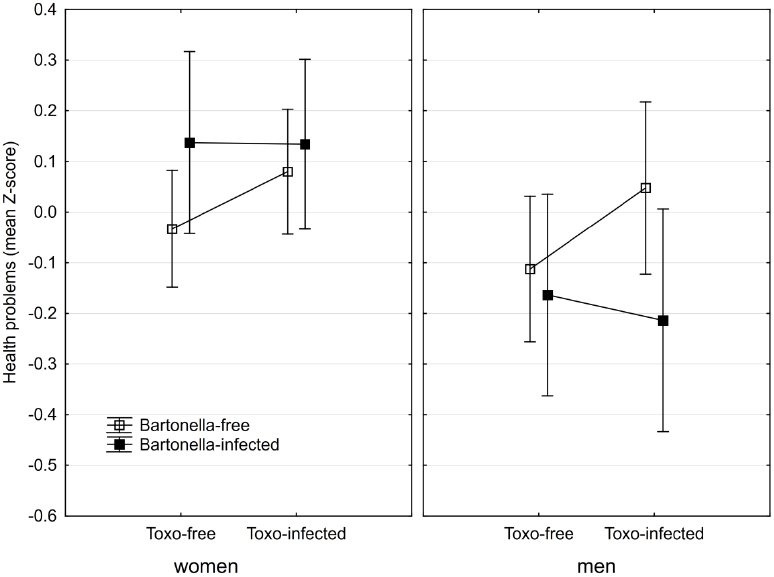
Health status of *Bartonella*- and *Toxoplasma*-seropositive subjects.

*Squares show the intensity of health problems computed for covariates at their means and spreads 95% confidence intervals.*

#### 3.2.4. Hypothesis 4

*The effects of Bartonella seropositivity on depression and health are stronger in Toxoplasma-seropositive subjects.*

The figures 1-3 show that there is no simple trend in the modulating effects of *Toxoplasma* seropositivity on the effects of *Bartonella* seropositivity. Moreover, we did not explicitly state, but we implicitly expected that the associations of *Bartonella* seropositivity with depression would be positive and with health would be negative. However, the associations of the *Bartonella* seropositivity with depression in *Toxoplasma*-seropositive men was negative – *Bartonella*-seropositive *Toxoplasma*-seropositive men had better mental health than *Bartonella*-seronegative *Toxoplasma*-seropositive men (Figs. 1, 2) and a similar paradoxical effect of the *Bartonella* seropositivity in the *Toxoplasma*-seropositive men was also observed on physical health (Fig. 3).

#### 3.2.5 Preregistered follow-up analyses

A MANCOVA analysis performed with six N-70 facets (except depression) as well as all seven N-70 subscales provided qualitatively the same result; therefore, only the result of the preregistered MANCOVA test with six subscales will be reported. The independent variables were sex, age, size of settlements where the responders spent their childhood, the *Bartonella* seropositivity, the *Toxoplasma* seropositivity, and the *Bartonella*-sex, *Bartonella*-*Toxoplasma*, and *Bartonella*-*Toxoplasma*-sex interactions. The only significant association observed was that of the *Bartonella*-*Toxoplasma*-sex interaction (p = 0.042, η^2^ = 0.045, Hotelling’s Trace method). The results of univariate ANCOVA analyses showed that this interaction was significant for the anxiety (p = 0.018, η^2^ = 0.033), obsession (p = 0.017, η^2^ = 0.034), hysteria (p = 0.008, η^2^ = 0.034), hypochondria (p = 0.048, η^2^ = 0.025), vegetative lability (p = 0.003, η^2^ = 0.049), and psychastenia (p = 0.010, η^2^ = 0.038); for the direction of these associations see Fig.1. Full models with all three 2-way and one 3-way interactions only showed a trend for the *Bartonella*-*Toxoplasma*-sex interaction (MANCOVA: p = 0.070, η^2^ = 0.048) and the effects of this interactions on obsession (p = 0.011, η^2^ = 0.027), hysteria (p = 0.011, η^2^ = 0.027), vegetative lability (p = 0.014, η^2^ = 0.025), and psychastenia (p = 0.023, η^2^ = 0.021). The results of nonparametric tests partial Kendall correlation tests performed separately for *Toxoplasma*-seropositive and *Toxoplasma*-seronegative male and female subjects are shown in Table 2.

**Table 2.** Correlation between *Bartonella* seropositivity and seven subscales of neuroticism in *Toxoplasma*-seronegative and *Toxoplasma*-seropositive subjects.

| N-70 subscale | *Toxoplasma*-<br>minus women<br>N = 81 |  | *Toxoplasma*-<br>plus women<br>N = 75 |  | *Toxoplasma*-<br>minus men<br>N = 54 |  | *Toxoplasma*-<br>plus men<br>N = 38 |  |
| --- | --- | --- | --- | --- | --- | --- | --- | --- |
|  | Tau | p | Tau | p | Tau | p | Tau | p |
| anxiety | -0.083 | 0.273 | -0.015 | 0.852 | 0.137 | 0.143 | -0.079 | 0.483 |
| depression | **-0.136** | 0.072 | -0.102 | 0.195 | 0.139 | 0.136 | **-0.210** | 0.063 |
| obsession | **-0.114** | 0.131 | 0.076 | 0.332 | 0.139 | 0.137 | **-0.210** | 0.064 |
| hysteria | **-0.207** | 0.006 | 0.060 | 0.449 | -0.010 | 0.918 | **-0.374** | 0.001 |
| hypochondria | 0.081 | 0.286 | 0.015 | 0.854 | 0.101 | 0.282 | **-0.184** | 0.105 |
| vegetative lability | **-0.129** | 0.089 | -0.107 | 0.175 | 0.191 | 0.042 | **-0.420** | <0.001 |
| psychastenia | **-0.161** | 0.033 | -0.028 | 0.723 | 0.104 | 0.265 | **-0.267** | 0.018 |

*Tau and p values of partial Kendall Tau correlations controlled for age. The results significant after the Benjamini-Hochberg procedure (false discovery rate = 0.2) are printed in bold.*

**Table 3.** Correlation between *Bartonella* seropositivity and health status in *Toxoplasma*-seronegative and *Toxoplasma*-seropositive subjects.

| Health problems | *Toxoplasma*-<br>minus women<br>N = 81 |  | *Toxoplasma*-<br>plus women<br>N = 77 |  | *Toxoplasma*-<br>minus men<br>N = 54 |  | *Toxoplasma*-<br>plus men<br>N = 38 |  |
| --- | --- | --- | --- | --- | --- | --- | --- | --- |
|  | Tau | p | Tau | p | Tau | p | Tau | p |
| allergic problems | **0.163\*** | 0.031 | 0.003 | 0.973 | 0.056 | 0.549 | -0.044 | 0.696 |
| dermatologic problems | **0.255\*** | 0.001 | 0.121 | 0.119 | **-0.254\*** | 0.007 | **-0.291\*** | 0.010 |
| cardiovascular problems | **0.298\*** | 0.000 | 0.075 | 0.335 | -0.123 | 0.189 | **-0.241\*** | 0.033 |
| digestive organs problems | 0.100 | 0.187 | 0.009 | 0.903 | -0.038 | 0.686 | -0.078 | 0.488 |
| metabolic problems | -0.008 | 0.919 | 0.039 | 0.619 | -0.127 | 0.175 | -0.220 | 0.051 |
| orthopedic problems | -0.023 | 0.762 | -0.166 | 0.032 | 0.040 | 0.672 | -0.044 | 0.700 |
| neurological problems | 0.077 | 0.312 | 0.114 | 0.142 | -0.215 | 0.022 | -0.032 | 0.780 |
| psychiatric problems | **0.189\*** | 0.013 | 0.125 | 0.113 | 0.156 | 0.096 | 0.075 | 0.506 |
| headache | -0.066 | 0.380 | 0.095 | 0.219 | -0.028 | 0.768 | -0.053 | 0.639 |
| other chronical pain | **0.201\*** | 0.008 | -0.076 | 0.327 | 0.124 | 0.186 | -0.132 | 0.242 |
| other chronical problems | **0.151\*** | 0.048 | 0.028 | 0.722 | 0.009 | 0.925 | -0.048 | 0.674 |
| tired | **0.180\*** | 0.018 | 0.028 | 0.720 | -0.023 | 0.803 | **-0.265\*** | 0.019 |
| tired after work | 0.122 | 0.107 | 0.003 | 0.973 | 0.122 | 0.192 | -0.145 | 0.201 |
| tired after travelling by train | 0.126 | 0.095 | -0.019 | 0.806 | -0.153 | 0.102 | -0.097 | 0.393 |
| viral infections | 0.114 | 0.133 | 0.020 | 0.797 | 0.118 | 0.208 | 0.131 | 0.246 |
| doctors' visits per year | **0.156\*** | 0.039 | -0.036 | 0.640 | 0.065 | 0.484 | 0.087 | 0.443 |
| antibiotics in last year | -0.014 | 0.853 | 0.144 | 0.063 | -0.058 | 0.537 | 0.034 | 0.761 |
| antibiotics in last 3 years | -0.063 | 0.404 | **0.294\*** | 0.000 | -0.094 | 0.316 | -0.214 | 0.058 |
| hospitals in last year | -0.071 | 0.348 | -0.160 | 0.039 | n.d. |  | n.d. |  |
| hospitals in last 3 years | -0.094 | 0.216 | -0.092 | 0.237 | -0.105 | 0.262 | -0.209 | 0.065 |

The table shows partial Kendall Tau (age controlled) and corresponding $p$ values.
Absence/presence of the specific antibodies was coded as 0/1, therefore, the positive Tau
means more serious or more frequent problem in the seropositive subjects. Results significant after the correction for multiple tests are labelled with asterisks. Correlation with number of hospitalizations in the last year (n.d.) could not be computed for men, due to a low number of hospitalized subjects.

To show which kinds of health problems were responsible for the association of bartonellosis with the health of our responders, we performed 20 ordinal regressions for 20 different health-related variables (rated on the 7-points scales) as the dependent variables and sex, age, size of settlements where the responders spent their childhood, *Bartonella* seropositivity, *Toxoplasma* seropositivity, and *Bartonella*-sex, *Bartonella*-*Toxoplasma*, and *Bartonella*-*Toxoplasma*-sex interactions as the independent variables. The result showed no effect of the *Bartonella*-*Toxoplasma*-sex interaction. At the same time it showed a significant effect of the *Bartonella*-*Toxoplasma* interaction on heart disorders (higher problems in the *Bartonella*-seropositive subjects, especially in those who were also *Toxoplasma*-seropositive, p= 0.038), recurrent pain (higher in *Bartonella*-seropositive and *Toxoplasma*-seronegative, lower in *Bartonella*-seropositive, *Toxoplasma*-seropositive, p = 0.026), and frequency of being tired (higher in *Bartonella*-seropositive and *Toxoplasma*-seronegative, lower in *Bartonella*-seropositive, *Toxoplasma*-seropositive, p = 0.049). The sex-*Bartonella*-interaction was significant for cardiovascular disorders (higher problems in the *Bartonella*-seropositive subjects, especially in women, p = 0.009), frequency of being tired (lower in *Bartonella*-seropositive men, p = 0.022), and frequency of taking antibiotics within the past three years (higher in *Bartonella*-seropositive women and lower in *Bartonella*-seropositive men, p = 0.023). The main effect of the *Bartonella* seropositivity was observed only for variable psychiatric problems, (more serious or frequent problems reported by the seropositive subjects of any sex, p = 0.032). No significant effect of *Toxoplasma* seropositivity was observed.

The BDI-II questionnaire consists of 21 questions. To show which problems are responsible for the observed associations of *Bartonella* seropositivity and depression, we computed partial Kendall correlations of *Bartonella* seropositivity with responses of subjects to these 21 questions, separately for *Toxoplasma*-seropositive and *Toxoplasma*-seronegative male and female subjects, Table 4). Only the negative correlation with problems with sleeping was significant for women after the correction for multiple tests. In contrast, the responses to nearly all items of the scale (except pessimism concerning future and problems with sleeping) correlated positively with the *Bartonella* seropositivity in *Toxoplasma*-seronegative men. In *Toxoplasma*-seropositive men, the responses to five questions, namely 7 - self-dislike, 11 - agitation, 12 – loss of interest, 13 – indecisiveness, 14 – worthlessness, and 15 – loss of energy correlated negatively with *Bartonella* seropositivity, i.e., the men infected with both *Bartonella* and *Toxoplasma* reported lower levels of depression than men infected only with *Toxoplasma*.

**Table 4.**
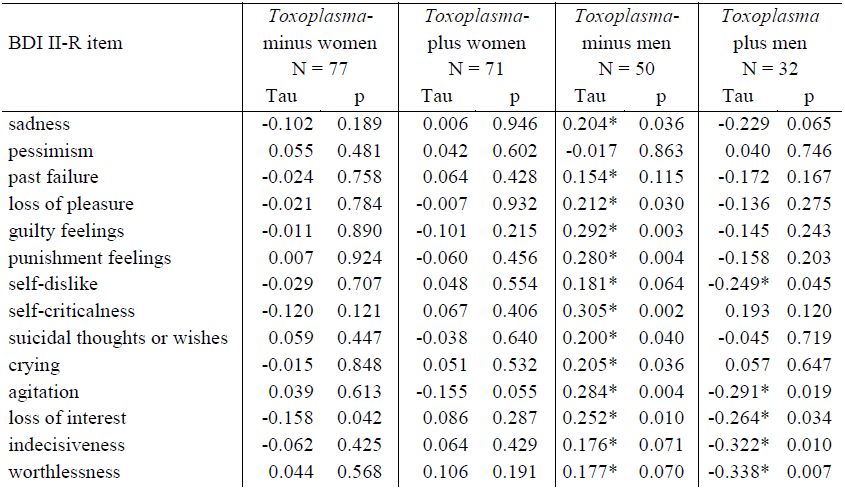
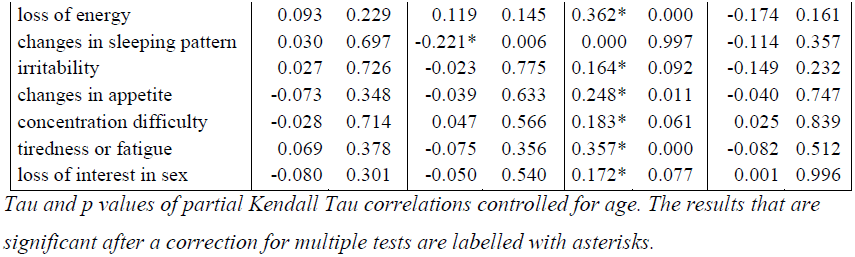
Correlation between *Bartonella* seropositivity and responses to BDI II questions in *Toxoplasma*-seronegative and *Toxoplasma*-seropositive subjects.

| BDI II-R item | *Toxoplasma*-<br>minus women<br>N = 77 |  | *Toxoplasma*-<br>plus women<br>N = 71 |  | *Toxoplasma*-<br>minus men<br>N = 50 |  | *Toxoplasma*-<br>plus men<br>N = 32 |  |
| --- | --- | --- | --- | --- | --- | --- | --- | --- |
|  | Tau | p | Tau | p | Tau | p | Tau | p |
| sadness | -0.102 | 0.189 | 0.006 | 0.946 | 0.204* | 0.036 | -0.229 | 0.065 |
| pessimism | 0.055 | 0.481 | 0.042 | 0.602 | -0.017 | 0.863 | 0.040 | 0.746 |
| past failure | -0.024 | 0.758 | 0.064 | 0.428 | 0.154* | 0.115 | -0.172 | 0.167 |
| loss of pleasure | -0.021 | 0.784 | -0.007 | 0.932 | 0.212* | 0.030 | -0.136 | 0.275 |
| guilty feelings | -0.011 | 0.890 | -0.101 | 0.215 | 0.292* | 0.003 | -0.145 | 0.243 |
| punishment feelings | 0.007 | 0.924 | -0.060 | 0.456 | 0.280* | 0.004 | -0.158 | 0.203 |
| self-dislike | -0.029 | 0.707 | 0.048 | 0.554 | 0.181* | 0.064 | -0.249* | 0.045 |
| self-criticalness | -0.120 | 0.121 | 0.067 | 0.406 | 0.305* | 0.002 | 0.193 | 0.120 |
| suicidal thoughts or wishes | 0.059 | 0.447 | -0.038 | 0.640 | 0.200* | 0.040 | -0.045 | 0.719 |
| crying | -0.015 | 0.848 | 0.051 | 0.532 | 0.205* | 0.036 | 0.057 | 0.647 |
| agitation | 0.039 | 0.613 | -0.155 | 0.055 | 0.284* | 0.004 | -0.291* | 0.019 |
| loss of interest | -0.158 | 0.042 | 0.086 | 0.287 | 0.252* | 0.010 | -0.264* | 0.034 |
| indecisiveness | -0.062 | 0.425 | 0.064 | 0.429 | 0.176* | 0.071 | -0.322* | 0.010 |
| worthlessness | 0.044 | 0.568 | 0.106 | 0.191 | 0.177* | 0.070 | -0.338* | 0.007 |
| loss of energy | 0.093 | 0.229 | 0.119 | 0.145 | 0.362* | 0.000 | -0.174 | 0.161 |
| changes in sleeping pattern | 0.030 | 0.697 | -0.221* | 0.006 | 0.000 | 0.997 | -0.114 | 0.357 |
| irritability | 0.027 | 0.726 | -0.023 | 0.775 | 0.164* | 0.092 | -0.149 | 0.232 |
| changes in appetite | -0.073 | 0.348 | -0.039 | 0.633 | 0.248* | 0.011 | -0.040 | 0.747 |
| concentration difficulty | -0.028 | 0.714 | 0.047 | 0.566 | 0.183* | 0.061 | 0.025 | 0.839 |
| tiredness or fatigue | 0.069 | 0.378 | -0.075 | 0.356 | 0.357* | 0.000 | -0.082 | 0.512 |
| loss of interest in sex | -0.080 | 0.301 | -0.050 | 0.540 | 0.172* | 0.077 | 0.001 | 0.996 |

*Tau and p values of partial Kendall Tau correlations controlled for age. The results that are significant after a correction for multiple tests are labelled with asterisks.*

### 3.3. Nonregistered exploratory analyses

To see whether the deteriorated health status played a role in the effect of *Bartonella*- *Toxoplasma*-sex association on depression, we also included health status as another covariate into the ANCOVA test (dependent variable: Beck depression score; independent variables: health score, *Bartonella, Toxoplasma*, age, sex, size of place of living in childhood, and the sex-*Bartonella, Toxoplasma*-*Bartonella* and sex-*Toxoplasma*-*Bartonella* interactions). The health score had a strong effect on the BDI-II depression score (p < 0.00001, η^2^ = 0.108). However, the strength of the association of depression with *Bartonella*-*Toxoplasma*-sex remained approximately the same (p = 0.043, η^2^ = 0.028) as was detected in the model without the health score (p = 0.066, η^2^ = 0.024). The same analysis for depression measured with N-70 could not be done due to the highly significant result of Levene’s test of equality of errors (F_7,240_ = 3.00, p = 0.005).

To see whether deteriorated health status played a role in the effect of *Bartonella*-*Toxoplasma*-sex association on the neuroticism estimated using N-70 inventory, we also included health status as another covariate into the MANCOVA test and following six ANCOVA tests (independent variables: *Bartonella*, age, sex, size of place of living in childhood, *Toxoplasma*, and the sex-*Bartonella, Toxoplasma*-*Bartonella* and sex-*Toxoplasma*-*Bartonella* interactions). The MANCOVA showed that health status had a strong effect on total neuroticism (p < 0.002, η^2^ = 0.086) and on five of six of it’s subscales, the anxiety (p = 0.004, η^2^ = 0.034), obsession (p < 0.0001, η^2^ = 0.077), hysteria (p = 0.104, η^2^ = 0.011), hypochondria (p = 0.012, η^2^ = 0.026), vegetative lability (p = 0.004, η^2^ = 0.034), and psychastenia (p = 0.034, η^2^ = 0.018). Again, the strength of association between neuroticism and *Bartonella*-*Toxoplasma*-sex was approximately the same as it was detected in the model without the covariate health – (MANCOVA: p = 0.043, η^2^ = 0.045, ANCOVAS: anxiety p = 0.023, η^2^ = 0.031, obsession p = 0.015, η^2^ = 0.035, hysteria p = 0.009, η^2^ = 0.039, hypochondria p = 0.060, η^2^ = 0.023, vegetative lability p = 0.003, η^2^ = 0.048, and psychastenia p = 0.011, η^2^ = 0.037).

Toxoplasmosis is known to have negative effects on the health of infected subjects. To check whether these effects can be also be detected in our dataset, we performed nonparametric analyses searching for the association between toxoplasmosis and health-related variables on the whole population (not on *Bartonella*-seronegative and *Bartonella* seropositive subpopulations). Analogically, we searched for the associations between bartonellosis and health-related variables on the whole population (not on *Toxoplasma*-seronegative and *Toxoplasma*-seropositive subpopulations). Table 5 shows that some effects of the infections can be also recognized by this simplified approach; however, some associations were obscured by the opposite-direction shifts in particular subpopulations.

**Table 5.**
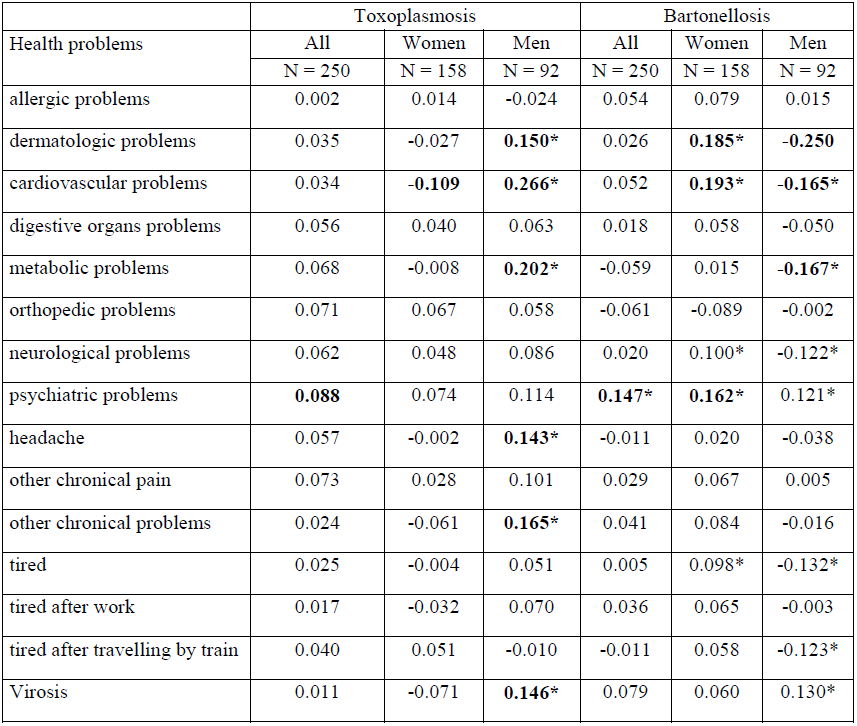
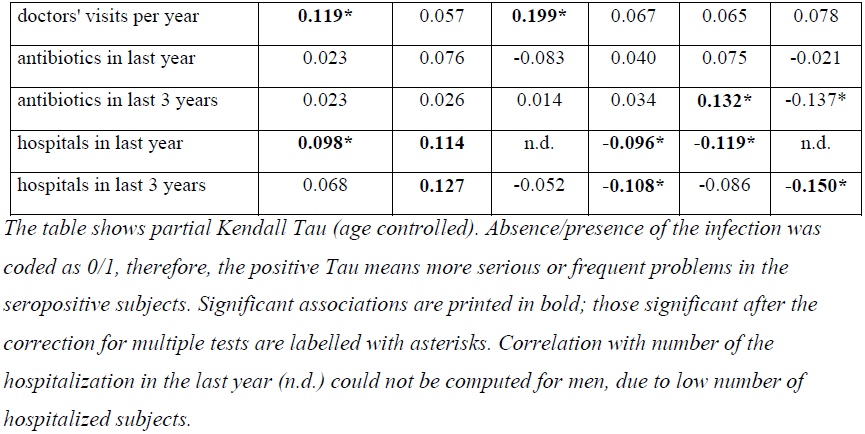
Effects of *Toxoplasma* and *Bartonella* seropositivity on health – one-way analyses.

Information about cat-related injuries was available for 213 subjects who responded to our email message in autumn of 2017. A Spearman correlation test showed no significant association between reported scratching by a cat and presence of anti-*Bartonella* antibodies. The results of a Spearman correlation of cat scratching with Beck depression score, N-70 depression, health, the presence of anti-*Bartonella* antibodies and presence of anti-*Toxoplasma* antibodies is shown in table 6. The relation between cat scratching, *Toxoplasma* seropositivity and Beck depression score is shown in Fig. 4.

**Table 6.** Association between animal-related injuries pathogens’ seropositivity and mental and physical health.

|  |  | *Bartonella*<br>IgG | *Toxoplasma*<br>IgG | Health<br>problems | Back<br>depression | N-70<br>depression |
| --- | --- | --- | --- | --- | --- | --- |
| dog biting | all | 0.002 | -0.01 | 0.001 | -0.068 | -0.030 |
|  | women | -0.046 | -0.036 | 0.007 | -0.061 | 0.12 |
|  | men | 0.084 | 0.043 | 0.021 | -0.068 | -0.075 |
| cat biting | all | -0.066 | 0.110 | 0.079 | **0.149\*** | 0.068 |
|  | women | -0.032 | **0.185\*** | 0.053 | **0.186\*** | 0.134 |
|  | men | 0.033 | -0.054 | 0.091 | 0.078 | 0.079 |
| cat scratching | all | 0.006 | 0.127* | 0.043 | **0.159\*** | 0.058 |
|  | women | -0.082 | **0.179\*** | 0.088 | 0.116 | 0.035 |
|  | men | 0.156 | 0.036 | -0.086 | **0.246\*** | -0.107 |

**Fig. 4.**
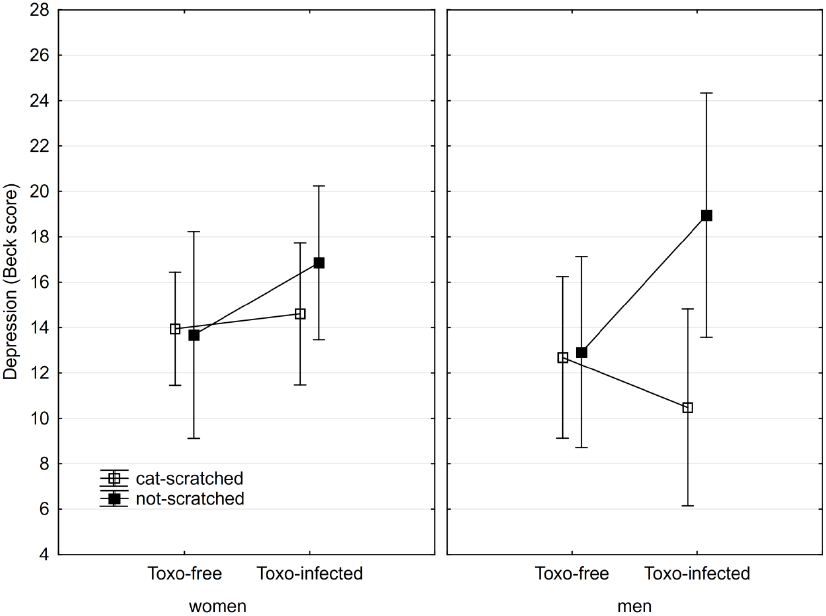
Association of cat scratching and *Toxoplasma* seropositivity with BDI-II depression score.

*Squares show depression computed for covariates at their means and spreads 95% confidence intervals.*

Twenty-eight (20.7 %) of 130 women and eight of (13.9 %) of 67 men reported using antidepressants and twenty-seven (20.6 %) of 131 women and five (7.6 %) of 66 men reported using anxiolytics. A strong correlation between taking antidepressants and anxiolytics was observed in both women (Spearman R = 0.681, p < 0.0001) and men (Spearman R = 0.452, p < 0.0001). The Beck depression score of the antidepressants-taking women was higher (Spearman R = 0.190, p = 0.030) while that of the antidepressants-taking men was lower (Spearman R = -0.223, p = 0.069) than the score of their non-drug-taking peers. No such correlations or trends were observed between taking antidepressants and the subscale of depression measured with N-70 (women: Spearman R = 0.053, p = 0.538, men: Spearman R = 0.019, p = 0.871) and no association was observed between scores of depression and taking anxiolytic drugs (Beck depression score, women: Spearman R = 0.142, p = 0.105, Beck depression score, men: Spearman R = -0.002, p = 0.986, N-70 depression score, women: Spearman R = 0.017, p = 0.849, N-70 depression score, men: Spearman R = 0.058, p = 0.624). The correlation between the Beck depression score and the N-70 depression score was non-significant neither for women (Spearman R = 0.120, p = 0.147), nor for men (Spearman R = 0.202, p = 0.069).

*The positive Spearman R always means positive association between the presence of anamnestic titres of anti-Bartonella or anti-Toxoplasma antibodies or with health, Beck depression, or N-70 depression (neuroticism) scores.*

## 4. Discussion

Our results suggest that the *Bartonella*-seropositive and *Bartonella*-seronegative subjects differ in mental and physical health. However, the observed pattern was different and more complex than we expected *a priori* on the basis of published data. The higher depression in *Bartonella*-seropositive subjects (hypotheses 1-2) was observed only in *Toxoplasma*-seronegative men and only when measured with the 21 questions of BDI-II test of depression (hypothesis 2), not with the10 items of the depression subscale from N-70 (hypothesis 1). The men seropositive for both *Bartonella* and *Toxoplasma* expressed lower level of depression than the subjects seropositive only for the *Bartonella*. This contrasted with the expectations of hypothesis 4, but agreed with recently published data, see below. No difference in the depression scores between *Bartonella*-seropositive and *Bartonella*-seronegative subjects was observed in women. Worse physical health, especially more frequent cardiovascular and dermatological problems, of *Bartonella*-seropositive subjects was observed only in *Toxoplasma*-seronegative women. Similarly, the *Bartonella*- and *Toxoplasma*-seropositive men reported better health, especially less frequent dermatological and cardiovascular problems, than the men seropositive only with *Bartonella* (which again contrasted with the expectations of hypothesis 4).

Our results seem to contradict the outcomes of two previous studies. The first study revealed a very strong association between having suffered a cat-related injury and being diagnosed with unipolar depression (Hanauer et al., 2013). The association was especially strong in women. The 9% prevalence of subjects diagnosed with major depression was observed among all 1.3 million patients of the University Hospital in Michigan. However, among those patients who were treated for a cat-related injury, 24 % of men and 48 % of women were also diagnosed with major depression within the period of 10 years (Hanauer et al., 2013). Based on these data we expected the positive association of depression and *Bartonella* seropositivity in both sexes, the stronger one in women. In reality, while our data show a relatively strong negative association between *Bartonella* seropositivity and physical health in women, the strong positive association with depression was observed only in (*Toxoplasma*-seronegative) men.

The second study (Flegr and Hodny, 2016) showed a positive association between suffering a cat-related injury, especially between being scratched by a cat, and reporting a major depression, and also between the cat-related injury and higher depression score measured with the BDI-II. Again, these associations were observed both in men and in women. However, after a careful inspection of the published data we realized that the odd phenomenon of the negative association between depression and suffering a cat scratching, the proxy for the *Bartonella* seropositivity, in *Toxoplasma*-seropositive men was also observed (but not commented) in the previous study (Flegr and Hodny, 2016). Figure 4 of that study shows that the Beck depression score for 15 non-scratched *Toxoplasma*-seropositive men was 18 (C.I._95_ = ±5), while this score for 31 scratched *Toxoplasma*-seropositive men was 14 (C.I._95_ = ±3.5). Corresponding scores for *Toxoplasma*-seronegative men were 10.2 (C.I._95_ = ±2.1) for 88 non-scratched men and 12.8 (C.I._95_ = ±2.4) for 90 scratched men. These scores are rather similar to those shown in Fig. 2 of the present study (means 18, 13.2, 10.5, and 16.8), despite the fact that scratching by a cat and the presence of anti-*Bartonella* antibodies was used as a proxy of the *Bartonella* seropositivity in the past and the present study, respectively.

The third study (Flegr and Vedralova, 2017) showed a significant correlation between reporting to be scratched by cat and thirteen of twenty-four studied mental health disorders, namely major depression, (OR = 2.97, CI_95_: 1.86–4.77), burn-out syndrome (OR = 1.88, CI_95_: 1.33– 2.66), panic disorder (OR = 2.18, CI_95_: 1.31–3.63), drug use disorder (OR = 3.3, CI_95_:1.45–7.53), antisocial personality disorder (OR = 2.02, CI_95_:1.1–3.7), sexual disorder (OR = 1.77, CI_95_: 1.06–2.97), obsessive compulsive disorder (OR = 1.63, CI_95_: 1.02–2.61), phobia (OR = 1.28, CI_95_: 1.0–1.65), borderline personality disorder (OR = 2.16, CI_95_: 1.0– 4.68), anxiety disorder (OR = 1.4, CI_95_: 0.98–2.0), bulimia & anorexia (OR = 1.94, CI_95_: 0.95–3.96), alcohol use disorder (OR = 1.61, CI_95_: 0.93–2.81), and other disorders (OR = 1.67, CI_95_: 0.89– 3.14) on a cohort of nearly nine thousand of internet users. The results of that study seem to agree with our results. Unfortunately, the study did not analyze men and women separately.

It must be reminded that the present study, in order to test the prediction of the hypothesis 4 (existence of *Toxoplasma*-*Bartonella* interaction), used a case-control study design with a similar number of *Toxoplasma*-seropositive and *Toxoplasma*-seronegative subjects while the earlier cohort studies analyzed data drawn from general populations of internet users. Typically, the *Toxoplasma*-seronegative subjects are much more common than those *Toxoplasma*-seropositive subjects (who seem to be protected against negative effects of *Bartonella*) in a general population, especially among young people. This explains why being injured by a cat, the proxy of *Bartonella* seropositivity, has been shown to be positively associated with the depression and with the probability of reporting diagnosed major depression in all three cohort studies.

Currently, we have no explanation for the negative association between the depression score and *Bartonella* seropositivity in *Toxoplasma*-seropositive men. It can be speculated that *Toxoplasma* seropositivity has in some respect a positive influence on certain facets of the mental health of *Bartonella*-seropositive subjects. *Toxoplasma* seropositivity does not increase (Sutterland et al., 2015) the risk of major depression, or even decreases such risk in men (Flegr, 2015; Flegr and Horacek, 2017). The *Toxoplasma* genome is known to contain two genes for tyrosine hydroxylases, the enzymes that catalyze the rate-limiting step in the synthesis of dopamine (Gaskell et al., 2009). Dopamine is synthesized in and nearby the cysts of the parasite in the brain tissue of the infected host (Prandovszky et al., 2011; Martin et al., 2015), but see also (Wang et al., 2015). This could positively influence the risk of schizophrenia and obsessive compulsive disorder. However, it could also decrease the risk of major depression and also depressiveness measured with BDI-II in members of a nonclinical population. Also, many studies showed a strong positive association between molecular markers of inflammation and depression (Danner et al., 2003; Penninx et al., 2003; Ford and Erlinger, 2004; Elovainio et al., 2009). It is possible that the increased concentration of IL-10, which is characteristic for *Toxoplasma*-infected hosts (Gaddi and Yap, 2007; Matowicka-Karna et al., 2009) (but see also (Kaňková et al., 2010a)), can decrease depression by its immunosuppressive and anti-inflammation activities (DeckertSchluter et al., 1997; Wilson et al., 2005). It is important in the context of the observed gender-toxoplasmosis-bartonellosis interaction that the negative correlation between the inflammation marker CRP (C-reactive protein) and depression was also reported to exist in men, but not in women; for a survey and discussion see (Vetter et al., 2013). It seems urgent to compare the level of CRP in *Toxoplasma*-seropositive and *Toxoplasma*-seronegative, *Bartonella*-seropositive subjects.

The alternative explanation of the observed pattern is that the men infected with both *Toxoplasma* and *Bartonella* could have such severe depressions that they must regularly use antidepressants and therefore report lower suffering from depressions. This explanation was supported by the fact that taking antidepressants correlated negatively with depressiveness measured by BDI-II in 67 male participants of the present study. In the *Toxoplasma*-seropositive subjects, both in men and women, cat scratching (in contrast to *Bartonella* seropositivity) had a positive, not negative, effect on the Beck depression score, compare the figures 2 and 4. This suggest that another factor, possibly another cat scratching-transmitted pathogen beyond *Bartonella henselae*, could be responsible for the observed positive association between the cat-related injuries and depression.

The strength of the statistical effect (η^2^) of *Bartonella* seropositivity increased rather than decreased when the effect of health was controlled. This suggests that the association between *Bartonella* seropositivity and depression is the result of some relatively specific effect, rather than being a non-specific side effect of the impaired health of *Bartonella* seropositive subjects.

The positive association between *Bartonella* seropositivity and depression (observed only in *Toxoplasma*-seronegative men) was much stronger when measured with the Beck inventory, which was used in the previous study (Flegr and Hodny, 2016), than with the N-70 questionnaire. We found no significant association between depressiveness measured with these two psychodiagnostic instruments. Moreover, our preliminary factor analysis of all 31 questions from these two questionnaires showed the existence of two distinct factors, the first one loaded exclusively with 21 questions from the Beck questionnaire and the second exclusively with 10 questions focusing on depression from the N-70 questionnaire (results not shown). Another factor analysis for just 70 questions of the N-70 questionnaire showed that this psychodiagnostic instrument measured only one factor, which is loaded with all ten questions focusing on psychastenia and seven of ten questions focusing on depression. These alarming results should be confirmed on a larger set of data; however, for the present, we can conclude that the published results concerning particular subscales of neuroticism measured with the 50 years old N-70 questionnaire should be interpreted, at least, with caution.

### Limits and strength of present study

The number of participants in the present study (250) was high enough for the reliable detection of main effects of infections and their 2-way interaction. However, for a reliable analysis of 3-way interaction, collecting a larger dataset would be highly desirable. It is also known that toxoplasmosis affects Rh-positive and Rh-negative subjects differently (Flegr et al., 2008; Novotná et al., 2008; Flegr et al., 2010; Kaňková et al., 2010b). For example, the positive association between toxoplasmosis and six of seven N-70 subscales of neuroticism has been recently observed only in Rh-negative women (Šebánková and Flegr, 2017). Again, to detect such a 3-way (and possibly even 4-way, bartonellosis-toxoplamosis-Rh-Sex) interaction, a much larger set of participants should be obtained and analyzed.

It is not possible to decide based on associations detected with observational studies what is the cause and what is the effect. *Toxoplasma* has been shown to affect hosts’ behavior by the experimental infection of laboratory animals (Berdoy et al., 2000; Hodková et al., 2007; Vyas et al., 2007; Webster, 2007); no such data, however, are available for *Bartonella*. The probability that depression or impaired health status would cause the presence of the anamnestic anti-*Bartonella* antibodies is rather low. However, it is possible that some unknown third factor, for example immunodeficiency, could cause both impaired mental health (or depression) and the infection by *Bartonella*.

## 5. Conclusions

At the present time, the effects of various symbionts on a phenotype of humans is the subject of a growing number of studies. Our results, however, suggest that rather than main effects of particular symbionts, the interaction of different species symbionts, parasites and mutualists, and their interactions with genotype and environment should be always studied. As a minimum, the effect of symbionts should be always studied separately on men and women as they usually react differently, often in an opposite way, to the infection. The opposite behavioral reaction to the *Toxoplasma* infection has been explained by the opposite reactions of men and women on the chronic stress that could be associated with life-long infection (Lindová et al., 2006; Lindová et al., 2010). However, the present results, namely the negative effect of *Bartonella* seropositivity on physical health and the positive effect of *Bartonella* seropositivity on the mental health of *Toxoplasma*-seronegative women, and the positive effect of *Bartonella* seropositivity on both physical and mental health in *Toxoplasma*-seropositive men, suggest that the stress-coping hypothesis alone cannot explain all observed phenomena and that the physiological state of men and women could really react differently to some specific environmental factors, including pathogen infections.

## Conflict of interest

The authors declare that the research was conducted in the absence of any commercial or financial relationships that could be construed as a potential conflict of interest.

## Funding sources

The work was supported by project UNCE 204004 (Charles University in Prague) and the Czech Science Foundation (Grant No. P303/16/20958).

### Acknowledgements

We would like to thank Charlie Lotterman for his help with the final version of the paper.

## Author contributions

JF designed the study and performed the statistical analyses. PB analyzed the sera. All authors participated on interpretation of results and writing the manuscript.

